# A Novel and Efficient Approach for Screening Cancer Cell Specific Monoclonal Antibodies

**DOI:** 10.1101/566398

**Authors:** Xin-Hui Pei

## Abstract

Cancer cell specific antibodies are pivotal tools in developing new immunotherapies for treating cancers. However, acquirement of cancer cell specific antibodies is time-consuming and often arduous. To circumvent such a barrier, we developed a novel antibody-screening method that can be used to efficiently produce cancer cell specific antibodies by an ‘antibody filter’ mechanism. First, we used normal human cells to perform the immunization in mice and collected the antisera. Second, we used human cancer cells together with the antisera against normal human cells to immunize another batch of mice. Theoretically, the antisera were able to neutralize the antigens from normal human cells, and therefore specific antigens only expressed in cancer cells could take advantage of the immunization. Third, we screened positive clones that are specific for cancer cells but not normal cells. Using this conceptual method, we successfully obtained 11 monoclonal antibodies that are specific for a human liver cancer cells line (HepG2) but not for a normal human liver cell line (HH). In addition, these clones failed to recognize other human cancer cells originated from different tissues, further highlighting the specificity. Collectively, we provide a novel and effective approach for screening cancer cell specific monoclonal antibodies, which may significantly facilitate the development of new anti-cancer therapeutics.

## 1 Introduction

Immunotherapy is one of the current research hotspots of cancer treatment program [1]. The fundamental principles of cancer immunotherapy include activating T cell recognition and cytotoxicity to cancer cell, stimulating the complement pathway by cancer-specific antibodies, and using targeted therapy by specific drugs[2]. Therefore, obtaining cancer-specific antibody is one of pivotal steps for developing therapeutic tools for cancer. Indeed, cancer cells are malignant hyperplasia cells which can escape the immunological surveillance. These cells can disguise their surface information as normal cells to prevent being identified and killed by the immune system [3,4]. That is, the high similarity of cancer cells to normal cells in surface antigens makes it a very difficult process to obtain cancer-specific antibody.

To obtain cancer-specific monoclonal antibodies is the pivotal and rate-limiting step for research and development of antibody-based cancer immunotherapy. There are two widely used strategies for cancer-specific monoclonal antibody screening: the first one is using cancer cells or their extracts to immunize mice and obtaining monoclonal antibodies followed by negative selection of control normal cells [5]. The second one is performing bioinformatics or biological analysis to identify novel cancer cell specific membrane antigen prior to immunization[6]. Unfortunately, both of strategies have their flaws. As mentioned above, the difference between cancer cell and normal cell at the molecular level is not obvious. Therefore, for the first method, a large amount of clones may often need to be screened for obtaining an antibody that can specifically recognize cancer cells[7]. Due to the inefficiency, this method was basically abandoned. The second method is successful in specific cancer cells, such as in pulmonary carcinoma[8]. But due to the heterogeneity of cancer cells, it is not affirmative that all of cancer cell populations express the same biomarker steadily. Therefore, the screened antibody can only target the cancer cell expressing the protein locally. According to a curative effect assessment of 62 new developed cancer medicines including monoclonal antibody drugs from 2003 to 2013, most of the drugs have undesired effects [9]. Methodological innovation is necessary for achieving multiple tools efficiently and conveniently to be applied for cocktail therapy.

Our new method in this study provides 11 liver hepatocellular carcinoma (HepG2)-specific monoclonal antibodies based on antibody *in vivo* negative selection principle (Fig. 1). These 11 antibodies have specific recognition to HepG2 with no cross-reactivity to both normal human hepatocyte cell (HH cells) and other kinds of cancer cells we tested. Briefly, we used whole normal cells, whole cell lysates from normal cells or membrane protein extract from normal cells as immunogens to immunize mice to obtain the antiserum against human normal liver cell. We then injected this antiserum containing anti-human normal hepatocyte antibodies to another normal non-immune mouse followed by immunization with cancer cell HepG2. Because this mouse already had anti-human hepatocyte antibodies, these excess antibodies can recognize the same antigen epitopes between normal hepatocyte and HepG2 and deliver them to macrophages or other phagocyte cells to degradation. Thus the antigens that can be recognized by anti-human hepatocyte serum can no longer cause the immune response of the mouse. And the left antigens of HepG2 which are not recognized by the serum will continue to activate the immune system and cause specific B cell proliferation and differentiation. Finally, we isolated spleen cells and executed the fusion procedure to obtained specific monoclonal antibodies. Since the anti-serum has neutralized the same constituents of HepG2 and HH cell as described above, these components do not irritate the immune response of the mouse, so that the mouse B-cell is only reacted to HepG2-specific antigen. We called this method of negative screening antigens *in vivo* with normal cell anti-serum as "antibody filters" (Fig. 1).

**Fig. 1.**
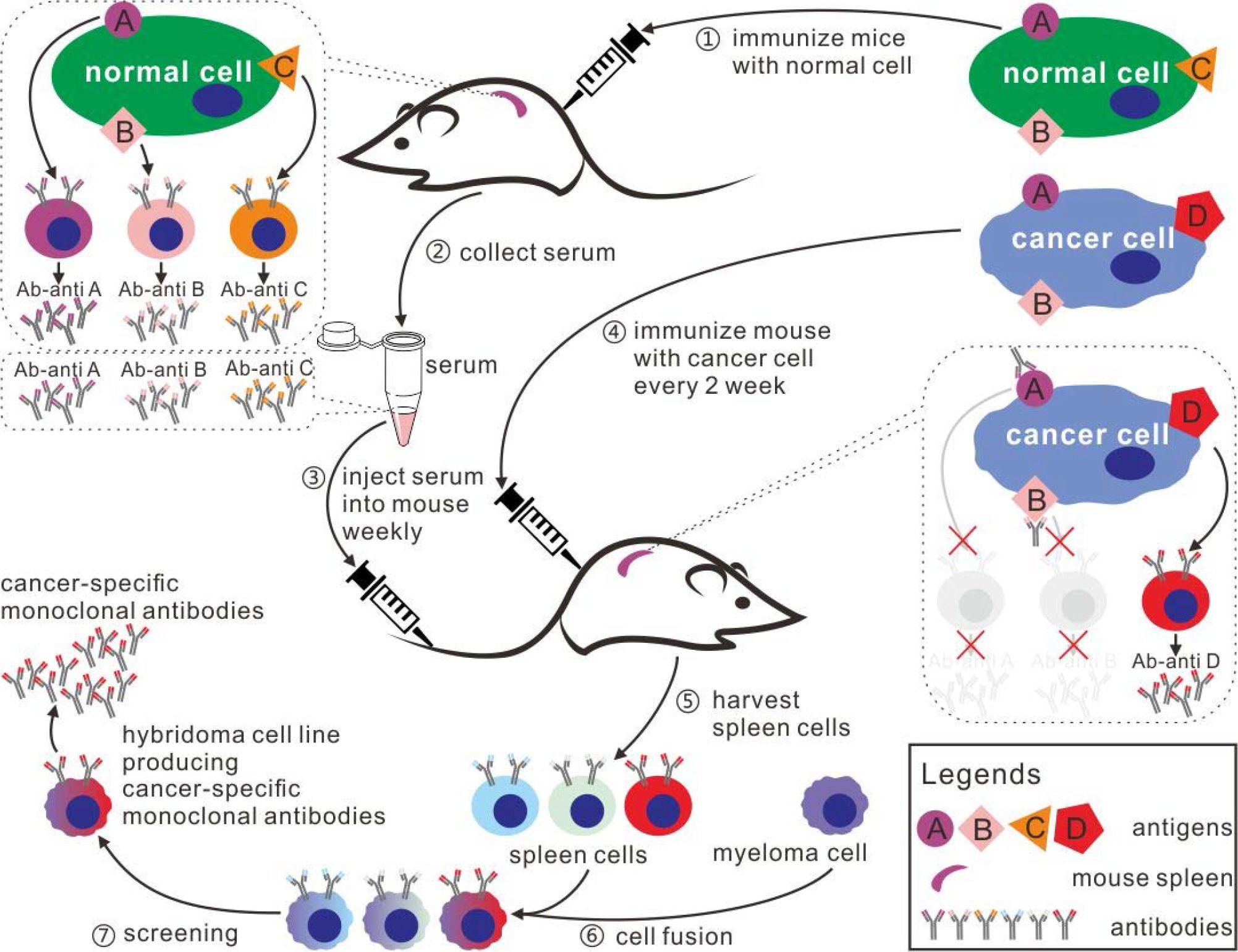
*An improved high-efficiency screening method for cancer-specific monoclonal antibodies.* Procedures: 1. immunize mice with normal cell; 2. Collect serum against normal cell; 3. Inject anti-normal serum into another mouse; meanwhile, 4. immunize this mouse with cancer cell every 2 weeks; 5. Harvest spleen cells of this mouse; 6. Proceed cell fusion; and 7. screen positive clones producing antibodies against cancer cell specifically.

## 2 Materials and methods

### 2.1 Cell culture

HepG2 cells and SP2/0 cells were purchased from the American Type Culture Collection (ATCC, Manassas, VA, USA). Dulbecco’s Modified Eagle Medium (DMEM), Fetal bovine serum (FBS) and streptomycin/penicillin were purchased from Invitrogen, Life Technologies, U.S.A. dimethyl sulfoxide (DMSO) and trypsin/EDTA and all other chemicals were purchased from Sigma Aldrich (Gillingham, Dorset, UK).

HepG2 cells were grown in monolayer cultures in Dulbecco’s Modified Eagle Medium (DMEM) supplemented with 10% fetal bovine serum, and 50 IU/ml penicillin and 50 μg/ml streptomycin. SP2/0 cells were grown in Dulbecco’s Modified Eagle Medium (DMEM) supplemented with 10% fetal bovine serum, and 50 IU/ml penicillin and 50 μg/ml streptomycin. Human Hepatocytes (Cat. No. 5200, ScienCell Research Laboratories, Inc.) were cultured in Hepatocyte Medium (Cat. No. 5201, ScienCell Research Laboratories, Inc.). Cells were maintained at 37°C in 5% CO2 with 95% humidity.

### 2.2 Primary cell immunization

Human Hepatocytes were primary cultured, collected, fixed with 1% paraformaldehyde (at room temperature for 0.5 hour), washed and stored at −20°C. A total of 50 female 6-week-old BALB/c mice were used for immunization. 100 million cells were intravenously injected per mouse. Mice were immunized once two weeks for a total of 3 times. Sera were collected 10 days after the last immunization.

### 2.3 Liver cancer cell immunization

HepG2 cells were cultured, collected, fixed with 1% paraformaldehyde (at room temperature for 0.5 hour), washed and stored at −20°C. A total of five 6-week-old BALB/c mice were intravenous injected with 0.3 ml anti-hepatocyte serum, followed with 1 million HEPG2 cells as immunogen. Mice were immunized once a week for a total of 3 times. After the last time of immunization, spleen cells were harvested and fused with SP2/0.

### 2.4 Screening of monoclonal antibody hybridoma cell lines

Separated spleen cells from immunized mice and SP 2/0 cells were mixed at the ratio of 5:1 and fused by PEG-1500. Fused cells were cultured in DMEM medium with supplement of HAT or HT. Positive monoclonal antibody clones were screened by ELISA assay in HepG2-coated 96-well plates. After three rounds of sub-cloning assay, the anti-HepG2 monoclonal antibody hybridoma cell lines were collected [10].

### 2.5 ELISA assay

Hepatocytes or HepG2 cells were seeded in 96-well plates. When the cell density reached 90-100%, wells were gently washed twice with PBS followed by PBS / 1% paraformaldehyde fixation. Plates were further rinsed with PBS buffer for 3 times. Culture supernatant of hybridoma or mice serum was diluted with PBS / 1% BSA and incubated with the coated well at 37°C for 1hr. HRP conjugated goat anti-mouse IgG secondary antibody diluted in PBS / 1% BSA were incubated after gentle wash at 37°C for 1hr. TMB substrates were used for color development in the dark for 30 minutes. Reactions were stopped by sulphuric acid. Absorbance of each well was read at 450 nm. Cut-off value was set as OD_450-blank_±0.500. Isotype of monoclonal antibodies derived from hybridoma supernatant was determined by Mouse Mab Isotyping Kit (Sino Biological) according to manufacturers’ instructions.

### 2.6 Antibody specificity

HepG2, 786-O, A375, DLD-1, ScaBER, A549, H441and WI-38 cells were seeded in 96-well plates. When the cell density reached 90-100%, wells were gently washed twice with PBS followed by PBS / 1% paraformaldehyde fixation. Plates were further rinsed with PBS buffer for 3 times. Culture supernatant from each hybridoma was incubated with the cell-coated well at 37°C for ELISA reaction. Culture supernatant from SP2/0 cell was used as blank. Absorbance of each well was read at 450 nm. OD_450_ of each well was divided by OD_450-blank_ as fold change.

## 3 Results

### 3.1 Immune serum against HH cell showed no observable difference in recognition between HH cell and HepG2

We immunized BALB/c mice with HH cells fixed with paraformaldehyde to obtain the anti-HH cell serum. Then we seeded HH cell and HepG2 into 96-well plate respectively, and made the serum gradient dilution. Serum was firstly diluted 1:1000 with PBS / 1% BSA, then executed 2-fold gradient dilution to the maximum dilution ratio of 2,048. In this study, we found that the ability of anti-HH cell serum to recognize HH cell and HepG2 was basically the same (Fig. 2).

**Fig. 2.**
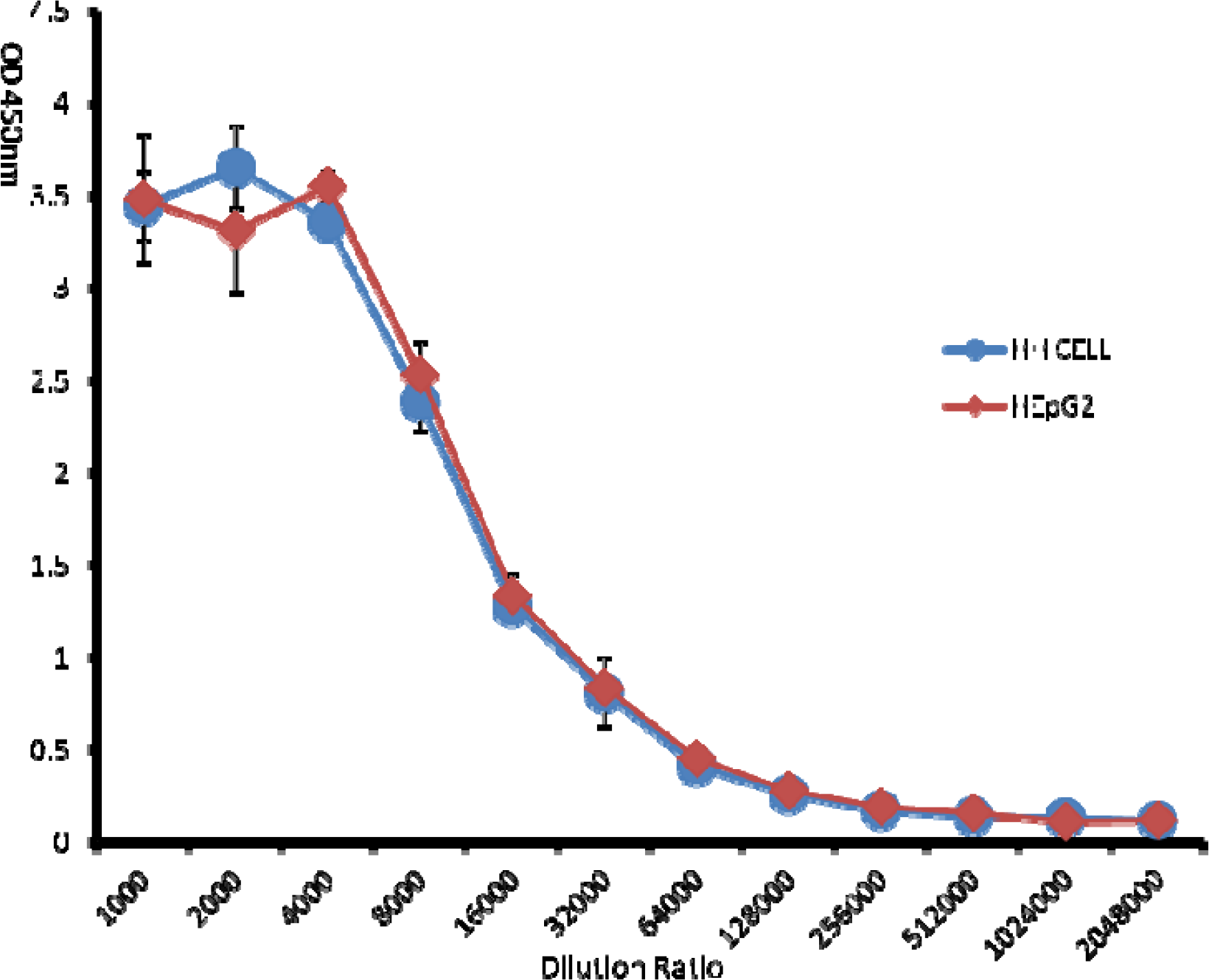
*Immune serum against normal human hepatocyte cell showed no observable difference to that of treated with hepatoma cell (HepG2).*

### 3.2 An improved high-efficiency strategy acquired 11 hybridoma cell lines producing monoclonal antibodies recognizing HepG2 cell specifically

We injected the anti-HH cell serum into 5 new BALB/c mice and immunized it with paraformaldehyde-fixed HepG2. During the immunization process, we injected the anti-HH cell serum to maintain the concentration of anti-HH cell serum in these mice. After the third immunization, spleen cells of these mice were taken and fused to SP2/0 cells. Finally, cells were screened with HepG2 cells, and 11 clones were obtained. We further test whether these antibodies can also recognize the HH cell by ELISA. We found that the 11 monoclonal antibodies were more clearly recognized for HepG2 rather than HH cell (Fig. 3).

**Fig. 3.**
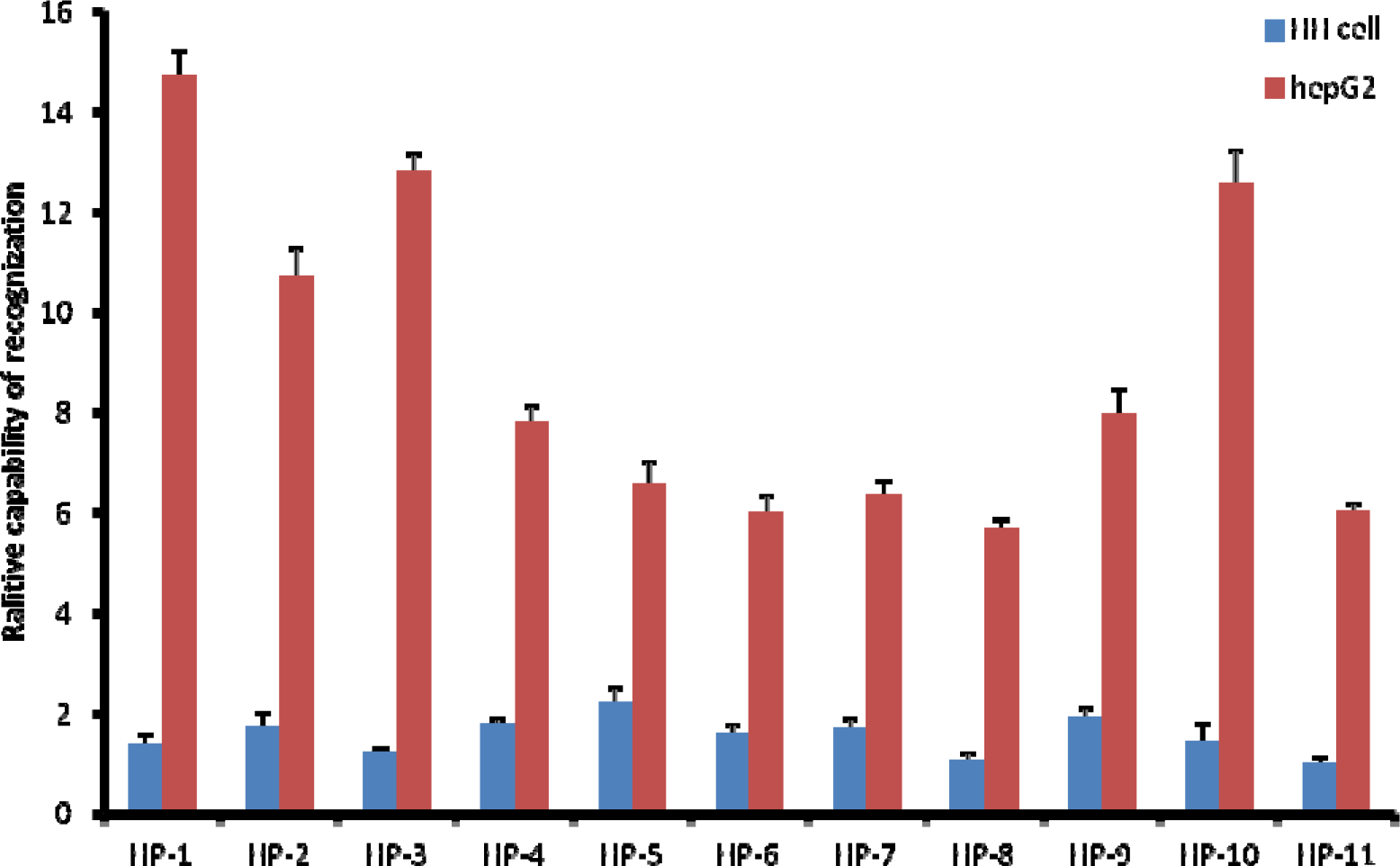
*An improved high-efficiency strategy acquired 11 hybridoma cell lines producing monoclonal antibodies recognizing HepG2 cell specifically.*

### 3.3 IgM accounted for up to 70% in those acquired monoclonal antibodies

Among these 11 monoclonal antibodies we obtained, HP1, HP2 and HP3 were IgG1 isotypes, while others were IgM isotypes (**data not show**). More interestingly, three IgG1 isotype (HP1, HP2, HP3) monoclonal antibodies showed stronger recognition ability to HepG2 than others (Fig. 3).

### 3.4 Monoclonal antibodies we produced have more specific recognition for liver cancer cells

We tested seven other sources of cancer cells or immortalized cells (786-O, A375, DLD-1, ScaBER, A549, H441and WI-38) respectively with these 11 antibodies. In our limited data, we found that these 11 antibodies showed a better ability to recognize HepG2 compared to other tissue-derived tumor cells or immortalized cells. (Fig. 4)

**Fig. 4.**
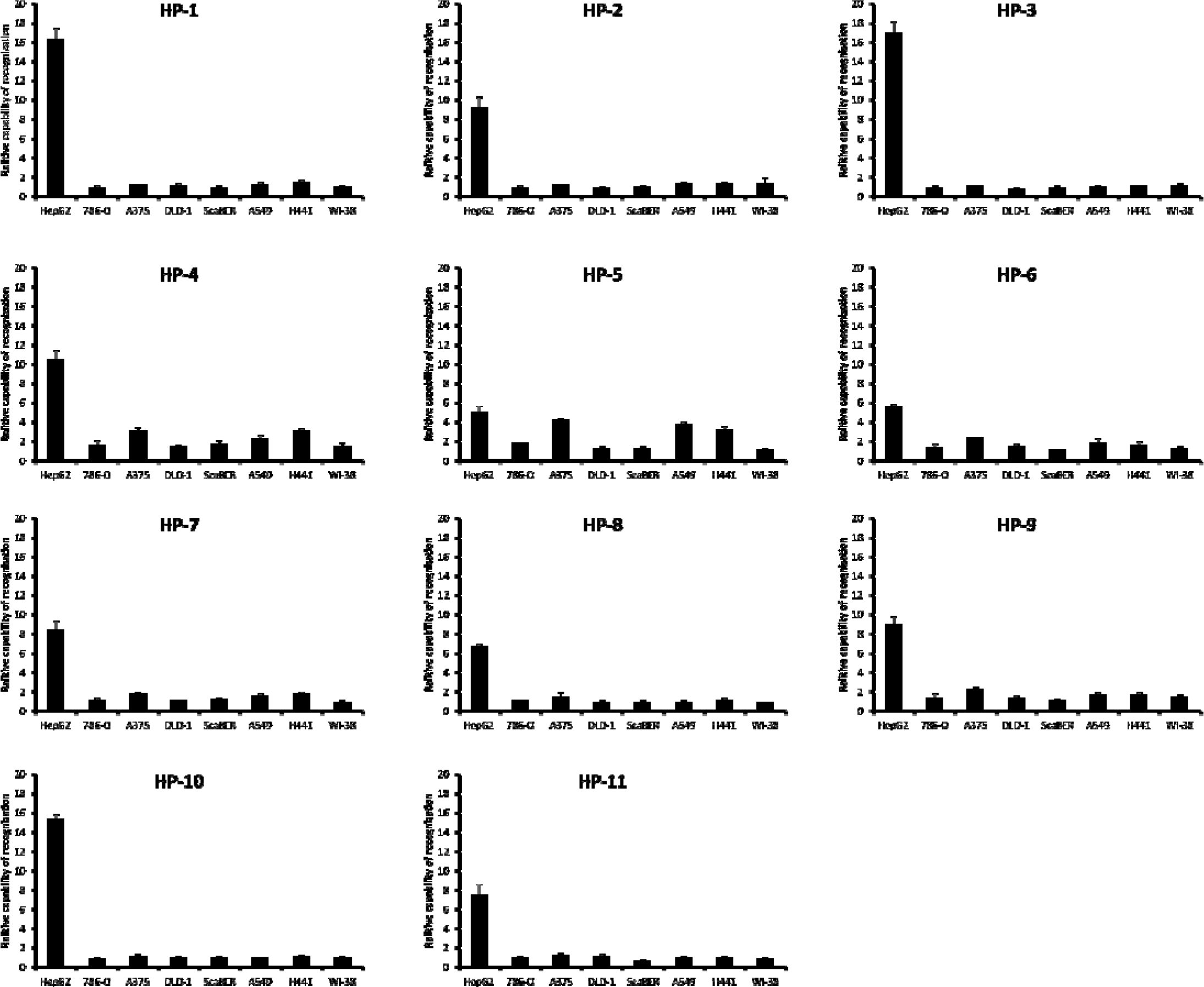
*Monoclonal antibodies we produced have more specificity for liver cancer cells.*

## 4 Discussion

The result of this study showed that after a series of dilution, abilities of anti-HH cell serum for recognizing HH cell and HepG2 cells are no difference, which means anti-HH cell serum can not distinguish HH cell and HepG2 cells. Otherwise, anti-HH cell serum would show a weaker recognizion capability to liver cancer cells than HH cells. These results indicate that the difference of membrane antigens between HH cell and HepG2 is very small (Fig. 2). This result may explain why antibodies obtained from traditional methods, for example immunizing mouse with hepG2 cell, can recognize normal liver cells unavoidably, which has brought great difficulty to screening. Namely, the traditional method of screening cancer-specific monoclonal antibodies with cancer cell immunization is a low efficient way [11,12].

Among these 11 hybridoma clones we obtained, we found that the isotype of 8 monoclonal antibodies was IgM, and it was generally believed that IgM isotype as low affinity antibody is produced because of failure to augment T-cell help with weak immunization or immunization deficient[13]. This result may indicate that HH cell is highly similar to HepG2 in the cell surface antigen. so that the anti-HH cell antibodies cleared most of the identical antigens of these two cell lines. The remained HepG2-specific antigens were difficult to stimulate the mouse immune response strongly. However, the eight IgM isotype antibodies are highly specific for HepG2 identification, which illustrates that the design of the method is reasonable (Fig. 3). In addition, we found that the three IgG1 isotype monoclonal antibodies showed a more significant recognition of HepG2 than that of IgM antobodies (Fig. 3), which indicates the existence of HepG2-specific antigens triggering sufficient immune responses. After the step of B cell clonal selection, these IgG1 isotype antibodies against HepG2 may have stronger identification abilities [14]. This result also suggests that if this method is further optimized, such as adjusting the ratio of the amount of immunized HepG2 cells and the amount of anti-HH cell serum injection, it is possible to obtain more IgG1, IgG2a or IgG2b isotypes with higher affinity. That is, the amount of HepG2 should be sufficient to achieve more secondary immune responses, and anti-HH cell serum should be sufficient to clear HH cell and HepG2 cross-antigens. We used the whole cell containing a lot of intracellular proteins to immunize the mouse, which may be the reason why a small number of antibodies were obtained and most of them were IgM isotypes. The next step, we intend to use membrane protein only, so theoretically we should obtain more anti-membrane protein specific antibodies [15].

To further explore the specificity of these monoclonal antibodies, we initially tested the ability of these antibodies binding to some cells. Clear cell renal cell carcinoma (786-O), melanoma cells (A375), colon cancer cells (DLD-1), bladder cancer cells (ScaBER), lung cancer cells (A549, H441) and human normal fibroblasts (WI-38) were tested for these 11 antibodies [16,17,18,19,20,21]. We found that these antibodies had weak affinity to these cells, and this result further demonstrates the specificity of these antibodies and suggests that there are significant differences of cell surface antigens in different types of cancer cells (Fig. 4). We also note that HP-5 has similar ability to identify some other cancer cells, which may suggest that the antigen of HP-5 corresponding to may be present on other types of cancer cell surfaces.

In this study, we obtained 11 antibodies by our invented screening technique. Although eight of them were IgM isotype, we found that these 11 antibodies had a better ability to recognize HepG2 comparing to HH cell or other cancer cells. It is important to note that the 11 antibodies were screened with HepG2 and were not screened with HH cells or other cells. In other words, the monoclonal antibody produced by this method does not need or rarely needs negative screening, which is impossible for the traditional method. This method undoubtedly greatly improved the screening effectiveness. Meanwhile, there is not necessary to know the specific antigens in our strategy, which is very helpful for those tumor cells that are difficult to find cell-specific antigens[22].

Furthermore, it also provides an easy method to identify differential protein profile in cells. For example, we can use these 11 antibodies to immunoprecipitate HepG2 lysate and obtain the proteins, which are likely specifically expressed in HepG2 relative to HH Cell. And then we can analyze them by mass spectrometry, which will undoubtedly help us to research the identified protein further[23]. This method can be used in different cells, such as final differentiation cells and stem cells. In addition to the cell membrane protein, the difference of intracellular proteins also can be identified by our method.

As mentioned above, for antibody immunotherapy of tumor, it is increasingly found that the treatment of single-type antibodies is not satisfactory. The tumor itself may have great diversities, and the single antibody treatment may exacerbate patient’s condition by screening out the more malignant cancer cells. So, more and more scientists believe that multiple antibodies, cocktail treatment, should be treated at the same time to minimize the escape of cancer cells[24]. However, in the traditional way, it is very difficult to obtain multiple specific monoclonal antibodies for a certain tumor at the same time. In our limited practice, we screened 11 specific antibodies, which may be an alternative method to obtain more tumor-specific antibodies. The best choice we can take is a variety of specific antibodies combination at the same time, which theoretically can significantly reduce the probability of tumor cell immune escape regardless of whether it is through the activation of complement pathway or antibody conjugated drugs to kill cancer cells.

In addition, our method can quickly obtain a type of cancer cell-specific antibody, which may be rapidly applied to the treatment of cancer. For example, we can obtain a specific patient’s tumor and its corresponding tissue cells, then quickly generate patient-specific tumor antibodies, and applied to tumor treatment in this way [25].

## Acknowledgments

*We thank Dr. Buqing Ye at Institute of Biophysics, Chinese Academy of Sciences for critical reading and discussion of the manuscript*.

## Conflict of interest

*The authors declare no financial or commercial conflict of interest*.

## Footnotes

* Several patents have been filed by Xin-hui Pei on the results of this paper.

## Abbreviations

HH cell: human hepatocyte cell
ATCC: American Type Culture Collection
DMEM: Dulbecco’s Modified Eagle Medium
FBS: Fetal bovine serum
DMSO: dimethyl sulfoxide

